# Regulation of the cell wall integrity pathway at the contact site between mating partners in yeast

**DOI:** 10.1101/2025.08.12.669915

**Authors:** Erin R. Curtis, Daniel J. Lew

## Abstract

The fungal cell wall is constantly remodeled to allow cell growth, but any holes in the cell wall would lead to catastrophic lysis. The “Cell Wall Integrity” pathway (CWI) detects cell wall defects and promotes cell wall thickening or repair to protect cell integrity. However, cell walls must be removed at contact sites between fusing cells during mating or mycelium formation. Here we show that in *Saccharomyces cerevisiae*, the CWI is downregulated specifically at the contact site between mating cells. A key component of the CWI, Pkc1, accumulated at polarity sites (shmoo tips) in cells exposed to mating pheromone, but not at contact sites. Pkc1 exclusion required a cell wall protein, Fig2, induced by pheromone. In mutants lacking Fig2, cell wall removal was delayed, blocked, or even reversed after transient fusion, leading to reduced mating. These results suggest that Fig2 designates the contact site as a “safe” spot to degrade the cell wall.

**eTOC:** Curtis and Lew show that the fungal “Cell Wall Integrity” repair pathway is silenced at contact sites between mating partners to allow cell wall degradation and fusion. They identify a cell wall protein needed to distinguish the contact site as a safe spot for wall removal.

## INTRODUCTION

Fungi are surrounded by a thin (100-200 nm) cell wall made of carbohydrate polymers and glycoproteins (Klis et al., 2002). In the hypoosmotic environments where fungi thrive, water influx leads to cell swelling and high turgor pressure. Fungal growth and proliferation requires localized weakening or thinning of the cell wall to enable directed expansion, but excessive weakening of the cell wall would lead to catastrophic cell lysis. The conserved fungal “Cell Wall Integrity” signaling pathway (CWI) protects cells from such lysis by detecting and responding to local defects that threaten wall integrity (Levin, 2005; Levin, 2011). In contrast, fungal cell-cell fusion requires complete removal of the cell wall at the fusion point (Clark-Cotton et al., 2022). How cells enable removal of the cell wall at this location but no other remains poorly understood.

The CWI detects cell wall defects through a set of transmembrane “sensors”, including Wsc1 and Mid2 in *Saccharomyces cerevisiae* (Lodder et al., 1999; Philip and Levin, 2001). The sensors have extensive extracellular domains and short intracellular domains that can recruit guanine nucleotide exchange factors (GEFs) to activate the GTPase Rho1 (Philip and Levin, 2001). Active Rho1 can bind and activate glucan synthases (Drgonová et al., 1996; Qadota et al., 1996) and also the conserved CWI kinase Pkc1 (Nonaka et al., 1995). Active Pkc1 phosphorylates multiple substrates, among them the kinase Bck1, which initiates a kinase cascade culminating in activation of the MAPK Slt2 (Nonaka et al., 1995). Active Slt2 (also called Mpk1) promotes transcriptional induction of genes encoding cell wall proteins (Jung and Levin, 1999). During polar growth of cells, local thinning of the cell wall at the growth site activates the CWI pathway to promote homeostatic re-thickening of the wall (Neeli-Venkata et al., 2021). Other conditions that lead to thinning of the cell wall, like hypo-osmotic shock (Davenport et al., 1995) and compressive stress (Mishra et al., 2017), also lead to CWI activation that is important for survival under stress.

In addition to promoting synthesis of cell wall proteins, CWI activation can lead to destabilization of cell polarity (the asymmetric accumulation of proteins and orientation of the cytoskeleton towards the growth site) following stress. This occurs following a sudden rise in temperature (Delley and Hall, 1999), applied compressive stress (Mishra et al., 2017), or laser-induced localized cell wall damage (Kono et al., 2012). In these circumstances, continued polar growth could lead to lysis and the transient reduction in polarization is thought to enable a corrective reset that allows for the repair of any cell wall thinning or damage before resuming polar growth.

Unlike the scenarios discussed above, there are occasions in the fungal life cycle when cell wall degradation is permitted. In particular, mating (Clark-Cotton et al., 2022), mycelium formation (Fischer and Glass, 2019) and generation of specialized structures like nematode traps (Nordbring-Hertz et al., 1989) involve cell-cell fusion that requires removal of the cell walls at the point of cell-cell contact (Clark-Cotton et al., 2022). This process is best understood in the mating reaction of *S. cerevisiae*. Haploid cells of each mating type respond to mating pheromones secreted by the opposite mating type and polarize growth towards each other (Sieber et al., 2023). At the contact site between cells, the two cell walls gradually thin until the two cells’ membranes touch and fuse (Gammie et al., 1998; Muriel et al., 2021). The CWI is activated by the polar growth that occurs in response to pheromone (Buehrer and Errede, 1997; Mishra et al., 2017) and mutation of CWI genes leads to lysis in response to pheromone (Hall and Rose, 2019; Ono et al., 1994). Thus, cell wall thinning during mating engages the CWI to protect cells from lysis. On the other hand, excessive CWI activation delays or blocks mating, presumably by preventing cell wall removal at the contact site (Hall and Rose, 2019; Philips and Herskowitz, 1997). How CWI activity is tuned to allow continued cell wall degradation despite cell wall thinning at the contact site is unknown.

Here we address how the CWI is spatiotemporally regulated during *S. cerevisiae* mating. We find that upstream CWI regulators accumulate at the contact site between mating partners, where cell walls are being degraded. However, Pkc1 shows a different pattern, with transient accumulation at multiple sites including locations adjacent to but not precisely at the contact site. These findings suggest that CWI action is spatially restricted to allow cell wall digestion only at the contact site. We further find that this restriction requires the pheromone-induced cell wall protein Fig2. In the absence of Fig2, CWI activity at the contact site delays, blocks, or even reverses cell wall degradation at that site. We suggest that Fig2 communicates to the CWI that the contact site is a safe spot to degrade the cell wall.

## RESULTS

### Localization of CWI components during yeast mating

In wild-type (WT) mating mixtures, cells of the opposite mating type undergo a search process in which polarity markers repeatedly relocate before stably clustering at the contact site between partner cells, marking “commitment” (Clark-Cotton et al., 2021; Henderson et al., 2019). After commitment, the cell walls at the contact site are thinned to allow for membrane apposition, and cell-cell fusion occurs about 25 min later (Fig. 1A,B). To investigate how CWI components behave during mating, we fluorescently tagged the CWI sensor Wsc1, the Rho1-GEF Rom 2, and the core CWI kinase Pkc1 (Fig. 1C). All of these components accumulate at sites of cell wall damage or polar growth (Kono et al., 2012; Lai et al., 2018; Neeli-Venkata et al., 2021). To assess protein behavior in the interval between commitment and fusion, we used two quantitative metrics. The first (Lai et al., 2018) uses the distribution of pixel intensities to report the degree of probe clustering (Fig. 1D). This metric is agnostic to the location of clustering.

**Figure 1:**
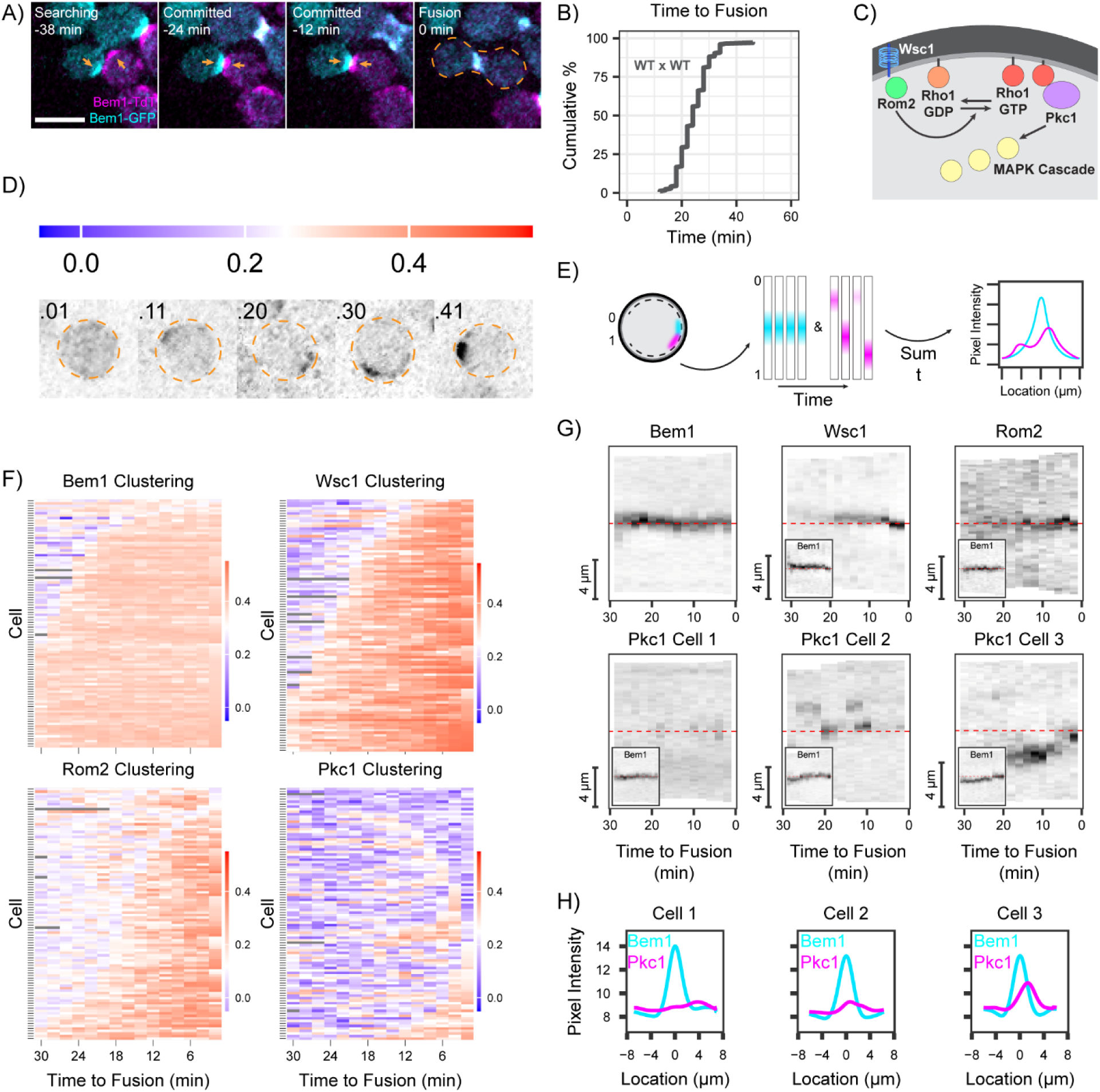
T**h**e **CWI in mating cells. (A)** Two-color time-lapse imaging of polarity probe Bem1 in mating cells. Maximum projection images (cyan, Bem1-GFP; magenta, Bem1-TdTomato) for selected timepoints. Scale bar, 5 μm. Strains: MAT**a** (DLY9069) x MAT*α* (DLY12944). Cell-cell fusion is indicated by orange dashed outline (0 min). Polarity sites prior to fusion indicated by orange arrows. **(B)** Cumulative distribution of the time from commitment (co-orientation of polarity sites) to fusion (n = 170 cells) for the same strains as in A. **(C)** Schematic depicting key components of the Cell Wall Integrity Pathway (CWI). **(D)** Schematic illustrating how different values of the clustering metric (color bar) relate to probe distribution. **(E)** Schematic depicting the pipeline from collecting linescans around the cortex of the cell, to generating kymographs, to compiling an intensity profile summed over time. **(F)** Clustering of fluorescently tagged probes (n = 100 cells) for polarity (Bem1-GFP, DLY24009 x DLY8156) or CWI components (Wsc1-GFP, DLY24254 x DLY8156; Rom2-mNeonGreen, DLY24325 x DLY8156; Pkc1-TdTomato, DLY24009 x DLY8156). Strains with tagged Bem1 and CWI probes were mated to an untagged partner so that fluorescence could be unambiguously assigned. Each row represents one cell. Color bar: degree of clustering ranges from highly clustered fluorescence (red) to uniform fluorescence (blue). **(G)** Kymographs of fluorescent signals from inverted maximum projection images of representative cells in the same mating crosses as in F. Linescans of the cell edges imaged at 2 min intervals for the 30 min before fusion, with the cell-cell contact site in the middle (red dashed line). Because Pkc1 behavior was more variable, three example cells are shown. For CWI probe kymographs, insets show Bem1 signal in the same cells. **(H)** Summed Bem1 and Pkc1 intensity profiles over time for the same cells displayed in G.

Cluster location was assessed by quantifying pixel intensities around the cell perimeter (linescans) (Fig. 1E). By tracing the same cell perimeter over time, the linescans generate a kymograph that displays the stability (or instability) of cluster locations (Fig. 1E). Summing these scans over time provides a graphical summary of the degree and location of probe clustering during the period of interest (Fig. 1E).

We focused on the 30 min prior to cell-cell fusion, when mating partners become committed and the intervening cell walls are degraded. During this period, the cell polarity probe Bem1 focused and remained tightly clustered at the cell-cell contact site (Fig. 1F,G). Consistent with previous findings in *S. pombe* (Neeli-Venkata et al., 2021), we found that Wsc1 became concentrated at mating contact sites shortly after polarity site alignment (Fig. 1F,G). Rom2 also became concentrated at contact sites shortly after Wsc1 (Fig. 1F,G), suggesting that Wsc1 becomes activated and recruits Rom2 to activate the CWI at mating contact sites, as it does during cell wall stress (Levin, 2005; Levin, 2011). Surprisingly, Pkc1 did not stably cluster at mating contact sites (Fig. 1F,G). Instead, Pkc1 was only sporadically clustered (Fig. 1F) at variable sites that often flanked the contact site between mating partners (Fig. 1G,H). Thus, cell wall thinning at the contact site appears to activate upstream CWI factors but not Pkc1, suggesting that Pkc1 is prevented from being activated at that site.

### Pkc1 behavior in shmoos differs from that in mating cells

The lack of Pkc1 accumulation at the contact site between mating partners was surprising, because previous work showed that exposure to mating pheromone led to activation of the downstream CWI kinase Slt2 (Buehrer and Errede, 1997). Consistent with that work, MAT**a** cells exposed to high levels of α-factor pheromone (in the absence of a partner) induced strong accumulation of Pkc1 at shmoo tips (Fig. 2A). The “shmoo” is a pear-shaped cell that develops when cells are exposed to high pheromone levels in the absence of a mating partner, due to arrest of the cell cycle in G1 and polarized growth. Pkc1 was often clustered at the shmoo tip along with the polarity marker Bem1 (Fig. 2A). Pkc1 clustering was stronger and more stable in shmoos than it was at mating contact sites (note different timescale as shmoos persist for much longer than contact sites)(Fig. 2B,C). Interestingly, kymographs showed that Bem1 clustering was broader and less stable at shmoo tips than at contact sites, and that Bem1 destabilization often coincided with strong Pkc1 recruitment to the shmoo tip (Fig. 2D. This frequent anti-correlation of Pkc1 and Bem1 signals (Fig. 2E) suggests that Pkc1 antagonizes tight Bem1 clustering at the shmoo tip. Over the course of a 3 h movie, Pkc1 clustering was much stronger at shmoo tips (Fig. 2F) than at mating contact sites (Fig. 1H).

**Figure 2:**
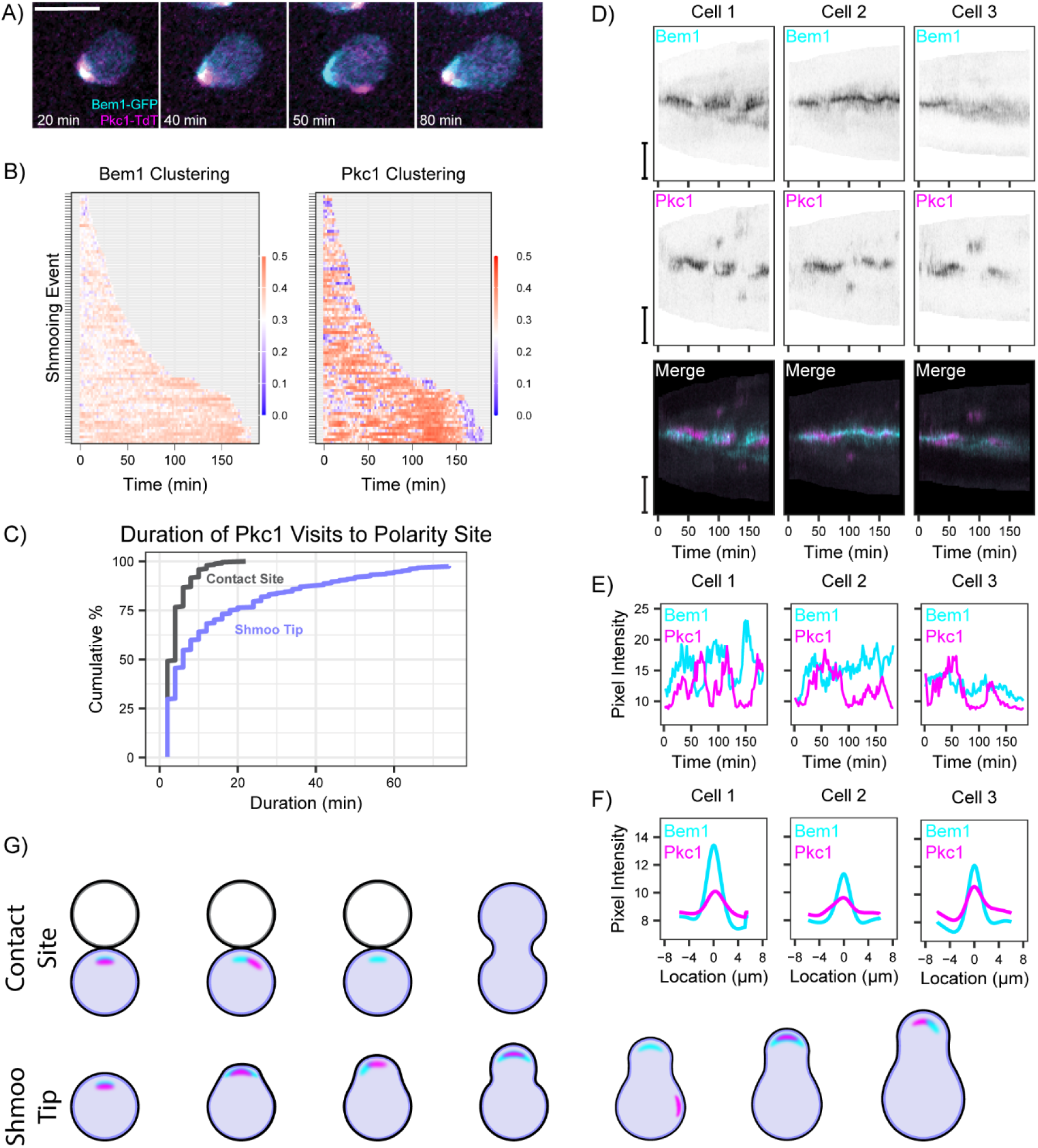
Pkc1 behavior in shmooing cells. **(A)** Two-color time-lapse imaging of Bem1-GFP (cyan) and Pkc1-TdTomato (magenta) in shmooing cell. Maximum projection images for selected timepoints. Scale bar, 5 μm. MAT**a** (DLY24009) cells treated with α-factor for the indicated times. **(B)** Clustering of Bem1-GFP and Pkc1-TdTomato probes in MAT**a** (DLY24009) cells treated with α-factor (n = 56 cells with 86 shmooing events). Shmooing events are periods when the Bem1 probe is continuously polarized. Note that the timescale is much longer than that shown in Fig. 1, and Pkc1 clustering values are higher than at contact sites (Fig. 1). Color bar: degree of clustering ranges from highly clustered fluorescence (red) to uniform fluorescence (blue). **(C)** Cumulative distribution of the duration of Pkc1 clustering in mating cells and shmoos of the same strain (DLY24009) (n = 100 mating cells with 240 Pkc1 visits, and n = 56 shmoos with 309 Pkc1 visits). p < 2.0×10^-12^ by Kolmogorov-Smirnov test. **(D)** Kymographs of fluorescent signals from maximum projection images of the same strains as in A-C. Linescans of the cell edges of representative shmoos were taken at 2 min intervals for 180 min, with the shmoo tip in the middle. Merge combines Bem1-GFP (cyan) and Pkc1-TdTomato (magenta) signal in the same cell. **(E)** Quantification of Bem1 and Pkc1 probe pixel intensities at the shmoo tips shown in D. **(F)** Summed Bem1 and Pkc1 intensity profiles over time for the same shmoos displayed in D,E. **(G)** Schematic summarizing Bem1 and Pkc1 behaviors at contact sites and shmoo tips.

Our findings indicate that Pkc1 behavior differs between shmoos and mating cells. As suggested by prior work, cell wall thinning at shmoo tips leads to strong Pkc1 recruitment, associated with partial destabilization of polarity. However, at the contact sites between mating cells, Pkc1 is rarely present, and polarity remains tightly focused (Fig. 2G). These observations suggest that some difference between shmoo tips and mating contact sites controls Pkc1 down-regulation at the contact sites.

### Role of mating agglutinins

A prominent difference between shmoo tips and contact sites is the engagement of mating agglutinins at contact sites. The agglutinins Sag1 (expressed only in MATα cells) and Aga2 (expressed only in MAT**a** cells) are cell wall proteins that bind to each other tightly (Zhao et al., 2001) and mediate adhesion between cells of opposite mating type (Cappellaro et al., 1991; Doi et al., 1989; Terrance and Lipke, 1981)(Fig. 3A). We speculated that Sag1-Aga2 engagement might distinguish contact sites from shmoo tips, providing a cue that indicates the presence of a contact site and down-regulates Pkc1 to allow cell wall removal at that site.

**Figure 3:**
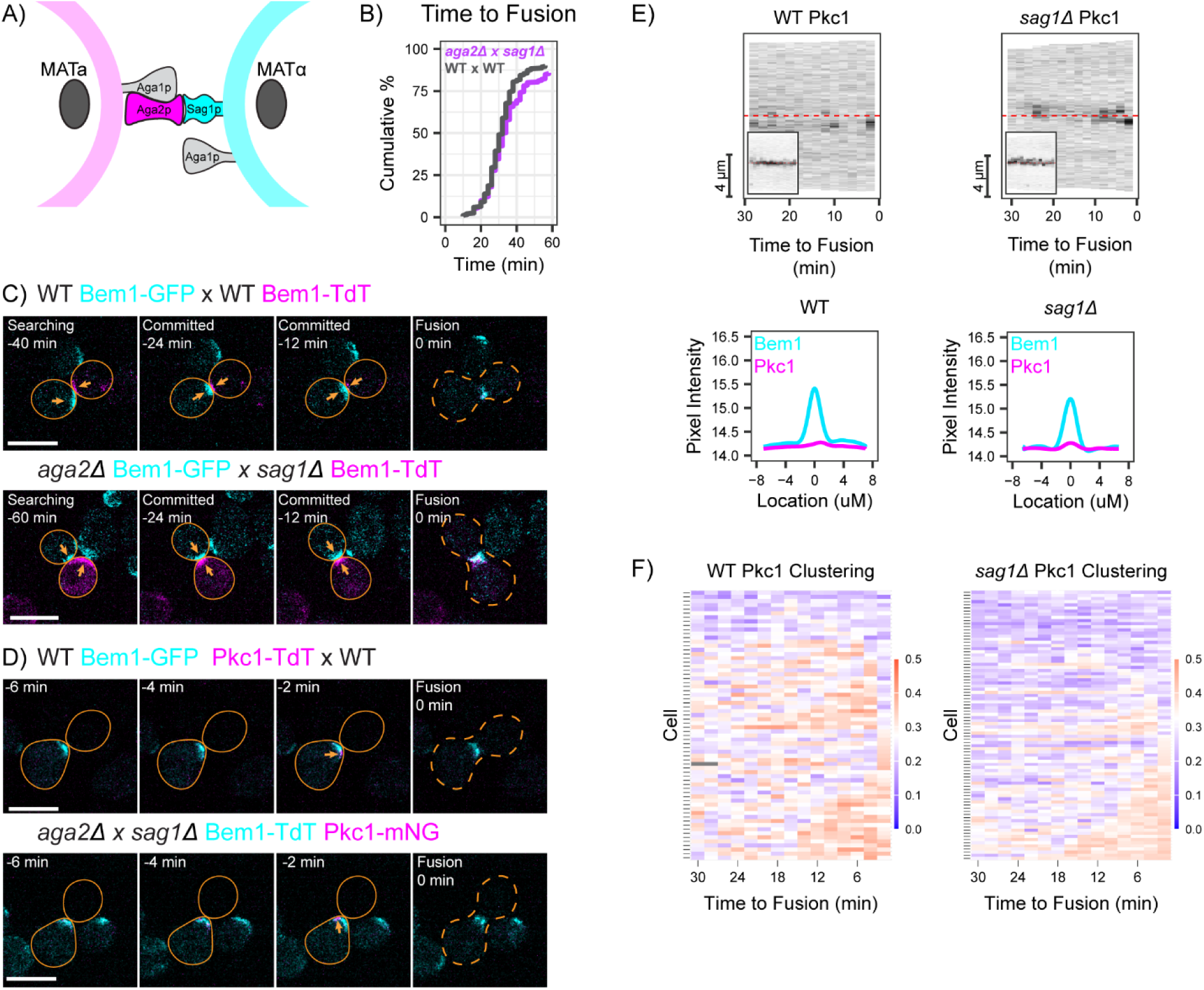
Mating is normal in the absence of agglutinins. **(A)** Schematic depicting the mating-specific agglutinins Aga2 and Sag1, which bind each other to adhere mating cells. Aga1 is expressed in both mating types, and anchors Aga2 to the cell wall in MAT**a** cells. **(B)** Cumulative distribution of the time from commitment (co-orientation of polarity sites) to fusion for the same strains as in A. WT: MAT**a** (DLY9069) x MAT*α* (DLY12944), n = 204 cells. Agglutinin-deficient: MAT**a** *aga2Δ* (DLY23477) x MAT*α sag1Δ* (DLY23471), n = 205 cells. p = 0.42 by Kolmogorov-Smirnov test. **(C)** Two-color time-lapse imaging of polarity probe Bem1 in mating cells. Maximum projection images (cyan, Bem1-GFP; magenta, Bem1-TdTomato) for selected timepoints. Scale bar, 5 μm. Crosses as in B. Cells are outlined with a solid orange outline and cell-cell fusion is indicated by orange dashed outline (0 min). Polarity sites prior to fusion indicated by orange arrows. **(D)** Pkc1 localization in mating cells. In these crosses, one partner expressing both Bem1-TdTomato (cyan) and Pkc1-mNeonGreen (magenta) probes was mixed with an opposite mating-type partner that lacked fluorescent probes. Maximum projection images for selected timepoints. Scale bar, 5 μm. WT: MAT**a** (DLY24009) x MAT*α* (DLY8156). Agglutinin-deficient: MAT**a** *aga2Δ* (DLY24657) x MAT*α sag1Δ* (DLY24655). Cells are outlined with a solid orange outline and cell-cell fusion is indicated by orange dashed outline (0 min). Pkc1 clusters indicated by orange arrows. **(E)** Kymographs of fluorescent signals from maximum projection images in the same mating crosses as in D. Linescans of the cell edges of representative cells were taken at 2 min intervals for the 30 min before fusion, with the cell-cell contact site in the middle. Insets: Bem1 signal in the same cells. **(F)** Clustering of fluorescently tagged Pkc1 (WT, n = 66; *sag1Δ* x *aga2Δ*, n = 86 cells) for the same crosses as in D.

Although Sag1 and Aga2 are important for mating in turbulent liquid media, they are not required for mating on solid media (Lipke et al., 1989; Roy et al., 1991). Consistent with that finding, we detected no mating defect when MATα *sag1* cells were mixed with MAT**a** *aga2* cells on agarose slabs. Mutant mating partners aligned their polarity sites and proceeded to fuse with similar timing to wild-type partners (Fig. 3B,C). Additionally, Pkc1 behavior in these mutant matings was similar to that in WT matings. In *sag1* x *aga2* crosses, Pkc1 was largely absent from the contact site in the minutes leading up until fusion, and as in wild-type crosses, it transiently clustered at a variety of sites (Fig. 3D-F). We conclude that mating agglutinins are not required to distinguish a contact site from a shmoo tip or to downregulate Pkc1 at contact sites.

### The cell wall protein Fig2 downregulates Pkc1 at contact sites

Previous studies identified several genes that were transcriptionally induced by pheromone, called α-*F*actor-*I*nduced *G*enes (*FIG*s)(Erdman et al., 1998). Among these, *FIG2* encodes a cell wall protein that accumulates at shmoo tips (Guo et al., 2000), and presumably at contact sites during mating. Cells lacking Fig2 displayed a somewhat confusing constellation of phenotypes (see Discussion), including hyperactivation of Slt2 in cells exposed to mating pheromone (Zhang et al., 2002). We speculated that Fig2 plays a role in down-regulating the CWI at contact sites between mating partners to identify the contact site as a safe place for cell wall removal.

In *fig2* x *fig2* crosses, Pkc1 was often concentrated at the contact site (Fig. 4A). In contrast, the polarity probe Bem1 was weaker and more mobile than in WT crosses (Fig. 4A). Using the clustering metric (Fig. 4B) confirmed that Pkc1 was more tightly clustered in mutant cells. This behavior resembled what we observed in shmoos (Fig. 2), suggesting that Fig2 plays a role in downregulating Pkc1, distinguishing the contact site from the shmoo tip. As discussed further below, *fig2* mutants also displayed prolonged commitment and mutant zygotes had narrower fusion bridges compared to wild-type (Fig. 4C, Fig. 5).

**Figure 4:**
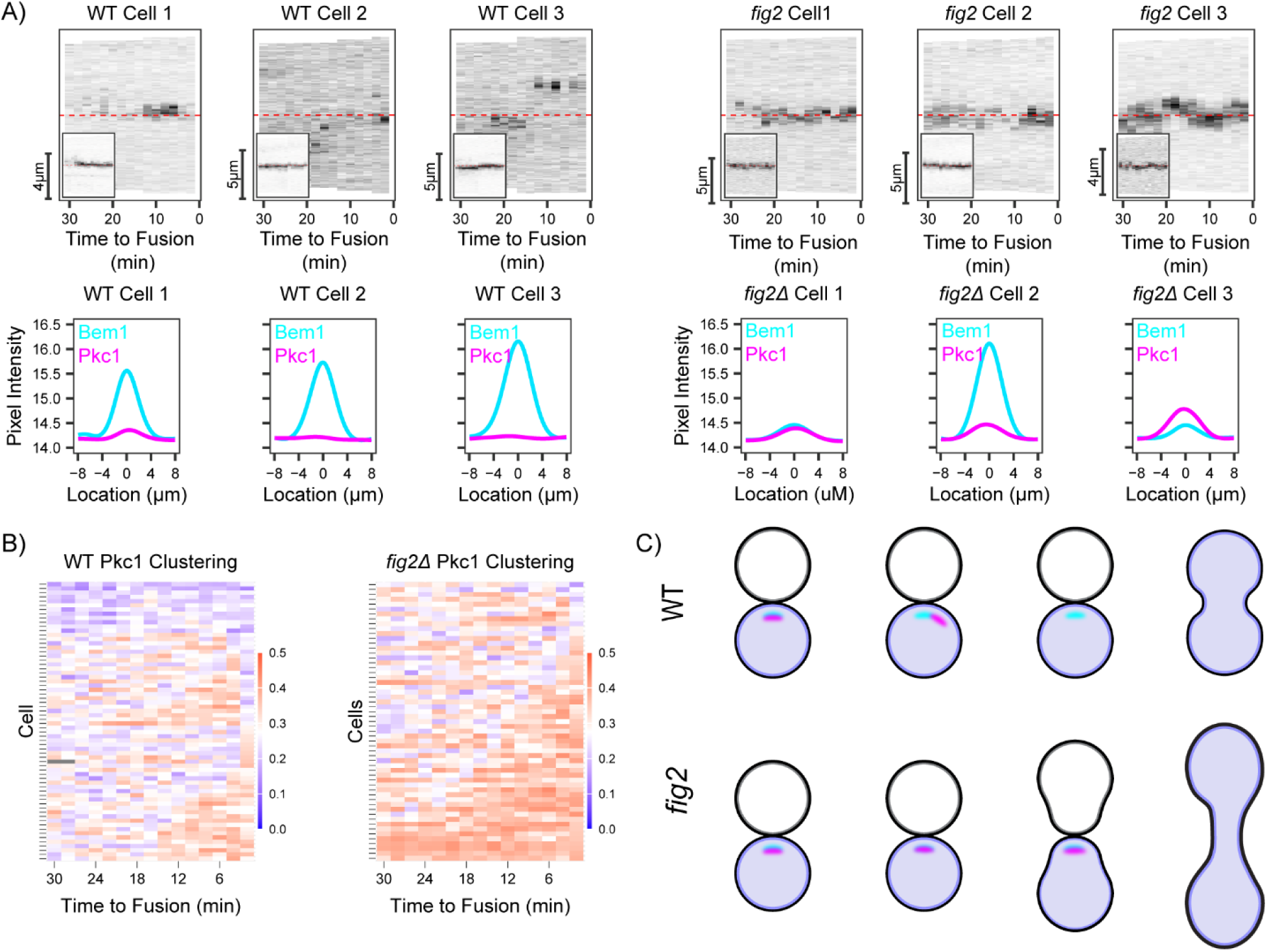
Fig2 is required for downregulation of Pkc1 at partner contact sites. **(A)** Kymographs and corresponding intensity profiles summed over time for Pkc1 and Bem1 fluorescent signals from WT (MAT**a**, DLY24009 x MAT*α*, DLY8156) and *fig2* (MAT**a** *fig2Δ*, DLY24546 x MAT*α fig2Δ*, DLY24551) crosses. **(B)** Clustering of fluorescently tagged Pkc1 (WT, n = 66; *fig2Δ* x *fig2Δ*, n = 57 cells) in the same crosses as in A. **(C)** Schematic highlighting the differential localization of Pkc1 in WT and *fig2* crosses.

**Figure 5:**
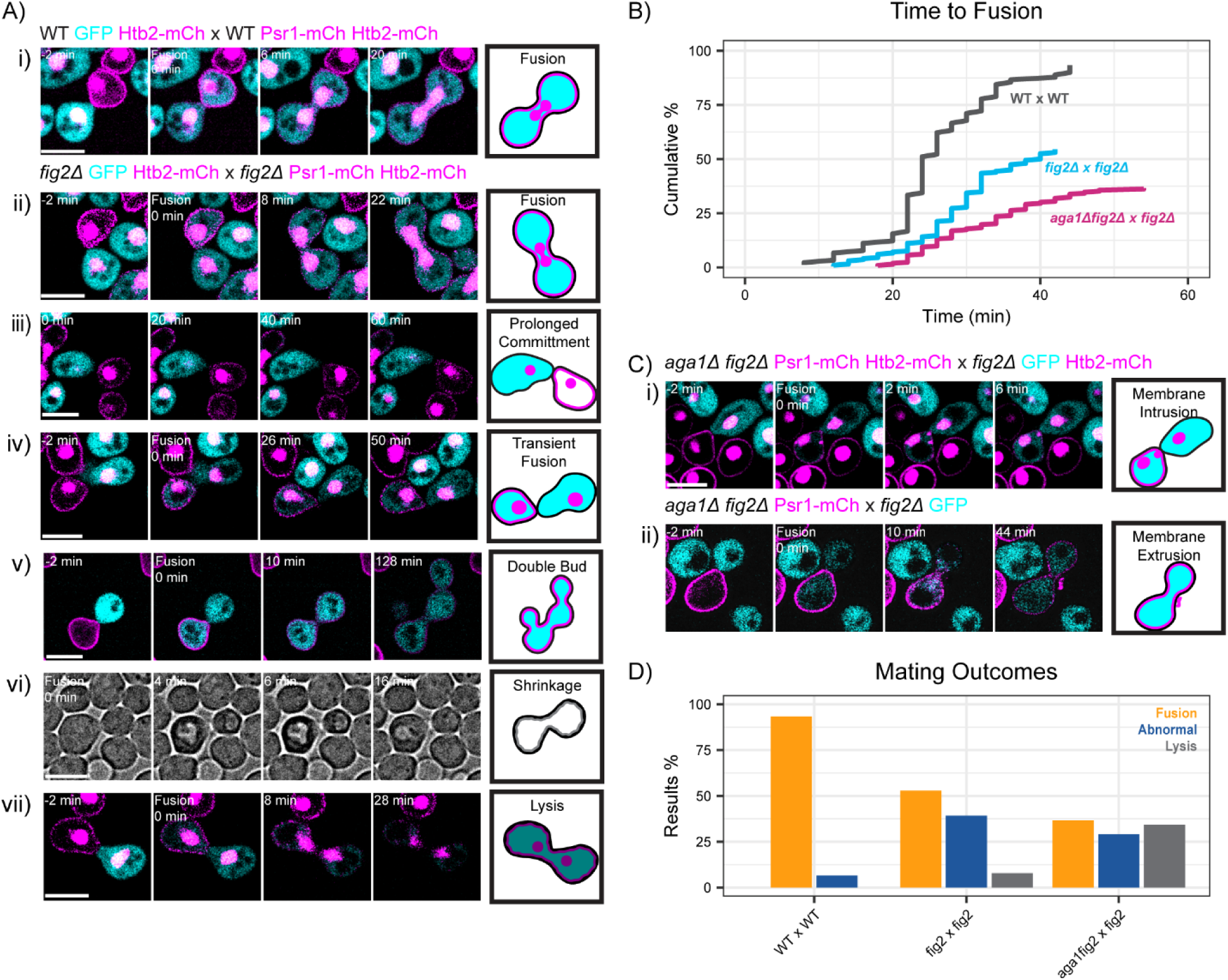
Mating defects in the absence of Fig2. **(A)** Two-color time-lapse imaging of cytoplasmic (GFP, cyan), membrane (Psr1-mCherry, magenta), and nuclear (Htb2-mCherry, magenta) probes in mating cells. In most panels, one partner expressed the cytoplasmic and nuclear probes while the other expressed the membrane and nuclear probes, enabling visualization of cytoplasmic mixing, membrane mixing, and nuclear fusion following mating. Maximum projection images for selected timepoints, with cell-cell fusion (cytoplasmic mixing) indicated at 0 min. Scale bar, 5 μm. WT: MAT**a** (DLY25323) x MAT*α* (DLY25324). *fig2*Δ x *fig2*Δ (i-iv and vi-vii): MAT**a** (DLY25321) x MATα (DLY25320). *fig2*Δ x *fig2*Δ (example v, strains lack nuclear probe): MAT**a** (DLY24672) x MATα (DLY26219). Each mutant example illustrates a different mating outcome, cartooned at right. In example vi) we show single-plane brightfield images to illustrate shrinking and re-swelling. See text for details. **(B)** Cumulative distribution of the time from commitment (co-orientation of polarity sites) to fusion for the same crosses as in A and C. WT x WT, n = 90 cells. *fig2*Δ x *fig2*Δ, n = 204 cells. *aga1*Δ *fig2*Δ x *fig2*Δ, n = 213 cells. For WT x WT vs *fig2*Δ x *fig2*Δ crosses p = 2.8×10^-9^ by Kolmogorov-Smirnov test. For WT x WT vs *fig2*Δ x *aga1*Δ *fig2*Δ crosses p < 2.2×10^-16^ by Kolmogorov-Smirnov test. For *fig2*Δ x *fig2*Δ vs *fig2*Δ x *aga1*Δ *fig2*Δ crosses p = 0.007 by Kolmogorov-Smirnov test. **(C)** Two-color time-lapse imaging as in A, for *fig2*Δ x *aga1*Δ *fig2*Δ cross: MAT**a** (DLY25319) x MATα (DLY25318). **(D)** Quantification of mating outcomes from the same crosses as in A-C. Outcomes are: fusion (regardless of timing: orange), attempted mating (prolonged commitment, transient fusion, partial lysis with or without fusion: blue), or lysis during mating (gray). For WT x WT vs *fig2*Δ x *fig2*Δ crosses p < 1.5×10^-10^ by Chi-squared test. For WT x WT vs *fig2*Δ x *aga1*Δ *fig2*Δ crosses p < 2.2×10^-16^ by Chi-squared test.

### Assorted mating defects in *fig2* x *fig2* crosses

The example *fig2* x *fig2* crosses illustrated in Fig. 4 were selected from those mating partners that mated successfully despite the Pkc1 misregulation, but many other mating pairs in the same cross failed to mate. In those cases we observed a diverse array of defects, including transient, reversible fusion attempts (Fig. 5). To investigate these events, we introduced probes to label the cytoplasm, plasma membrane, and nuclei. WT mating cells show fusion (mixing of cytoplasmic probe) followed by mixing of the plasma membrane probe and then karyogamy (fusion of the two nuclei)(Fig. 5Ai). A similar sequence occurred in about half of the mutant mating pairs, although fusion was delayed compared to WT (Fig. 5Aii, B). Other cells showed a prolonged commitment that did not result in fusion within the duration of the movie (Fig. 5Aiii). During the prolonged commitment, partners sometimes fused, as judged by mixing of the cytoplasmic probe, but then failed to mix the membrane or nuclear probes (Fig. 5Aiv). We interpret this to reflect a transient, reversible fusion in which cell walls are regenerated. In some cases, mating partners appeared to fuse but then went on to form separate buds (Fig. 5Av), behaving as if they were separate cells. In the bright-field images of transient fusion, some cells appeared to shrink and then re-swell (Fig. 5Avi), which suggests that transient fusion can be accompanied by a partial loss of cytoplasm, perhaps via transient cell wall gaps. In some cells, fusion was followed by irreversible lysis (Fig. 5Avii). Quantification of the various *fig2* mating defects indicated that about half of the mating pairs succeeded in fusing (albeit after variable delays) while the other half either failed to remove the intervening cell walls, or restored them after transient fusion, or lysed (Fig. 5B,D).

Fig2 and a related cell wall protein, Aga1, are thought to be paralogs created from a whole-genome duplication that occurred in the lineage leading to *S. cerevisiae* (Byrne and Wolfe, 2005). Aga1, like Fig2, is expressed in both mating types (Roy et al., 1991), but it has a MAT**a**-specific role in tethering the agglutinin Aga2 to the cell wall (Fig. 3A). In addition, work from the Erdman lab suggested that the Aga1 expressed in MATα cells has a second role that overlaps with that of Fig2 (Huang et al., 2009). Thus, we also examined matings in which both partners lacked Fig2 and the MATα partner also lacked Aga1 (this cross retains Aga1-Aga2/Sag1 agglutination: see Fig. 3A). This cross showed slightly more severe mating defects that were qualitatively similar to those of the *fig2* x *fig2* cross (Fig. 5B), including instances where fusion was accompanied by the sudden appearance intracellular (Fig. 5Ci) or extracellular (Fig. 5Cii) bodies labeled by our plasma membrane probe. These instances might represent partial and reversible lysis, with intrusion or extrusion of pieces of the plasma membrane.

The diverse array of phenotypes described above might all reflect a similar underlying cause: Pkc1 dysregulation. In the *fig2* x *fig2* cross, Pkc1 is activated at the contact site (Fig. 4). Local CWI activation may delay or prevent removal of the intervening cell walls, leading to prolonged commitment where mating partners are aligned but have not yet fused (Fig. 5Aiii). Similarly, CWI activation at the contact site may promote resynthesis of cell wall even after/during an initial fusion, leading to the observed transient fusion (Fig. 5Aiv). And focusing of CWI activity at the contact site may interfere with cell wall repair in adjacent regions, leading to partial (Fig. 5Avi, vii & viii) or complete (Fig. 5Av) lysis.

### Effect of the combined absence of Fig2 and agglutinins

Previous work indicated that *fig2* mutants and agglutinin mutants display synthetic mating defects (Guo et al., 2000). To assess how mating fails in such cells, we imaged mating pairs lacking both Fig2 and Aga1 in both mating types. Without Aga1, the MAT**a** agglutinin Aga2 is not anchored to the cell wall and agglutination fails despite the continued presence of Sag1 on the MATα partner (see Fig. 3A)(Roy et al., 1991). In these *aga1 fig2* x *aga1 fig2* crosses, polarity sites oriented towards each other for prolonged times but fusion did not occur (e.g. Fig. 6Ai maintained commitment for 70 min), suggesting that degradation of the intervening cell walls was not effective. At some point, one of the committed cells rotated (Fig. 6Ai 78 min), perhaps pushed by the partner. Cells that rotated often re-established polarity site co-orientation at the new contact site (Fig. 6Aii). In the absence of cell rotation, partners either remained committed throughout the movie (Fig. 6Aiii) or the polarity sites became destabilized (Fig. 6Aiv). These outcomes are diagrammed in Fig. 6E: we did not observe any successful mating events in this cross.

**Figure 6:**
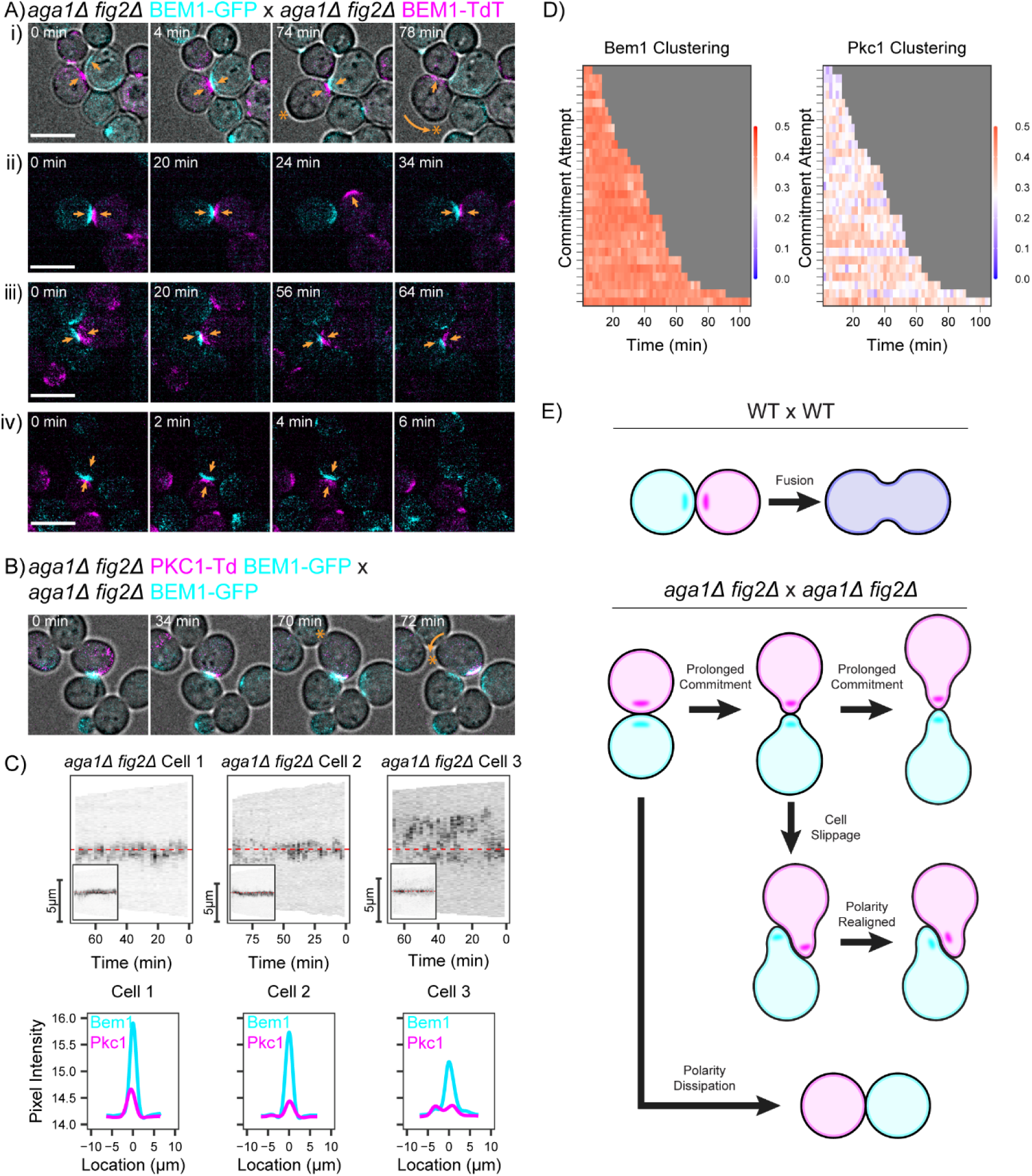
Mating defects in the absence of Fig2 and agglutinins. (A) Two-color time-lapse imaging of polarity probe Bem1 in mating mixes of cells lacking Fig2 and agglutinin engagement (*aga1*Δ mutants fail to attach Aga2 so there is no Aga2-Sag1 attachment despite the continued presence of Sag1). Cells of opposite mating type express differently tagged Bem1 (cyan, Bem1-GFP; magenta, Bem1-TdTomato). Maximum projection images for selected timepoints. Scale bar, 5 μm. Strains: MAT**a** *aga1*Δ *fig2*Δ (DLY23441) x MATα *aga1*Δ *fig2*Δ (DLY23438). Orange arrows indicate polarity sites. As these cells did not fuse, time 0 is arbitrary for each example. In the top example, brightfield images are merged with corresponding fluorescent images to make it easier to appreciate that cells slip and rotate (curved orange arrow indicates movement of region denoted by asterisk). See text for details. (B) Pkc1 localization in mating mixes of cells lacking Fig2 and agglutinin engagement. In these crosses, one partner expressing both Bem1-GFP (cyan) and Pkc1-TdTomato (magenta) probes was mixed with an opposite mating-type partner expressing only Bem1-GFP. Brightfield images are merged with corresponding fluorescent images to make it easier to appreciate that cells slip and rotate (curved orange arrow indicates movement of region denoted by asterisk). Maximum projection images for selected timepoints. Scale bar, 5 μm. Strains: MAT**a** *aga1*Δ *fig2*Δ (DLY24084) x MATα *aga1*Δ *fig2*Δ (DLY23438). **(C)** Kymographs and corresponding intensity profiles summed over time for Pkc1 and Bem1 fluorescent signals from the same crosses as in B. **(D)** Clustering of fluorescently tagged Pkc1 (n = 26 cells and n = 58 commitment attempts) for the same crosses as in B. **(E)** Schematic depicting fusion (WT) or failed attempt (mutant) outcomes in *aga1*Δ *fig2*Δ x *aga1*Δ *fig2*Δ crosses.

In similar crosses with a Pkc1 probe, we observed strong and prolonged recruitment of Pkc1 to the contact sites (Fig. 6B-D). These cells resembled the shmoos lacking a partner (Fig. 2). Thus, in the combined absence of Fig2 and agglutinins, it appears that cells fail to perceive the difference between a shmoo tip and a contact site. By failing to downregulate Pkc1 at that site, we propose that CWI activation prevents removal of the intervening cell walls, precluding mating.

## DISCUSSION

Here we report that the CWI is down-regulated at contact sites between yeast mating partners. Such down-regulation appears to be important to facilitate cell wall degradation at that site, allowing fusion between partners to yield diploid zygotes. Further, we show that the cell wall protein Fig2 is required for CWI down-regulation at the contact site. We suggest that Fig2 is a key component of the system that distinguishes the contact site from a shmoo tip. These findings, combined with those of previous studies in this area, leave several fascinating questions for future investigation, as discussed below.

### What is the mechanism of CWI down-regulation at contact sites?

CWI activation involves sensing of cell wall defects by transmembrane sensors (predominantly Wsc1 and Mid2), which then recruit Rho1-GEFs including Rom2 to activate Rho1 (Lodder et al., 1999; Philip and Levin, 2001). Active Rho1 then recruits and activates Pkc1 to promote cell wall repair (Nonaka et al., 1995). We found that Wsc1 and Rom2 both accumulated at contact sites between mating partners, but Pkc1 did not. These results are consistent with several possible mechanisms: (i) Rom2 is recruited but not active at contact sites. This would imply that the sensors not only recruit but also regulate the activity of downstream Rho1-GEFs. (ii) The recruited Rom2 does activate Rho1, but at the contact site a Rho1-GAP reverses that activation. This would imply that there is a location-specific means of regulating the GAP. (iii) Rho1 is indeed activated by upstream CWI factors at mating contact sites, but Rho1-GTP fails to recruit Pkc1 to that site (unlike other sites). Whatever the detailed mechanism(s) of CWI regulation, our findings suggest that there are location-specific influences on Pkc1 activation.

### How does Pkc1 locally counteract cell wall thinning?

The best-understood mechanism by which active Pkc1 counteracts cell wall thinning is the activation of a MAPK cascade leading to transcriptional induction of genes encoding cell wall proteins and cell wall synthases (Jung and Levin, 1999). However, this mechanism of action is global rather than local, in that Pkc1 activation anywhere on the cortex should lead to similar downstream effects. Instead, our findings suggest that when Pkc1 does localize to the contact site (in *fig2* mutants), it interferes with cell wall degradation at that site, whereas Pkc1 activation elsewhere is not problematic. Indeed, Pkc1 activation at other sites may help to explain how the CWI acts to counteract lysis during mating. These findings suggest that Pkc1 has additional local targets beyond the MAPK/transcriptional pathway that enable cell wall repair or rethickening specifically at the site of Pkc1 activation. The nature and functions of these targets remains to be determined.

One likely target of local Pkc1 action is the cell polarity machinery. Consistent with previous studies (Hall and Rose, 2019; Smith et al., 2017), we observed differences in polarity probe (Bem1) behavior between shmoo tips and mating contact sites. At contact sites, the lack of Pkc1 was associated with tightly polarized Bem1. At shmoo tips, strong Pkc1 recruitment was associated with weaker and less stable Bem1 localization, and episodes of Bem1 dispersal were temporally correlated with increased Pkc1 accumulation (Fig. 2). This is consistent with the notion that local Pkc1 activity weakens polarity. One possible Pkc1 target is the pheromone response scaffold protein Ste5, which is phosphorylated and inactivated by Pkc1 (Lee et al., 2020; van Drogen et al., 2019).

### How do mating cells distinguish a contact site from a shmoo tip?

As exposure to even saturating pheromone levels does not induce yeast cells to degrade the cell wall when not in contact with a partner, there must be a signal that differentiates a contact site from a shmoo tip. Perhaps the simplest possibility is that the difference is purely based on the contact-site geometry (Huberman and Murray, 2014). In this view, shmoo tips and contact sites need not differ in any physiological regard: both orient the actin cytoskeleton to secrete cell wall hydrolases at the polarity site, but these hydrolases do not suffice to thin the cell wall of the secreting cell. Only when two cells, each secreting hydrolases, have juxtaposed polarity sites does the local level of secreted hydrolase activity rise to the level required to thin the intervening walls (Huberman and Murray, 2014). While this model is elegantly simple, the situation appears to be more complex, because effective cell wall degradation requires a tighter focusing of the polarity machinery and actin cytoskeleton that only occurs following cell-cell contact (Dudin et al., 2015; Smith et al., 2017).

In the fission yeast *Schizosaccharomyces pombe*, polarity site focusing can be induced in the absence of a partner by engineering cells to secrete the pheromone that they normally receive from a partner (Dudin et al., 2016). Such partner-less focusing leads to wall degradation at the shmoo tip and hence lysis. These observations suggested that in the presence of a mating partner, the trigger for wall degradation is exposure to a spatially focused pheromone gradient (Dudin et al., 2016). In *S. cerevisiae*, other findings suggested that the trigger for wall degradation is contact-induced flattening of plasma membrane curvature (Smith et al., 2017).

Our findings suggest that in *S. cerevisiae*, cell-cell contact leads to local down-regulation of the CWI at the contact site, and that this allows cell wall degradation to proceed. Consistent with that view, attenuation of CWI activity by mutation of the sensor Mid2 results in polarity site focusing and lysis even in the absence of a partner (Hall and Rose, 2019). It is possible that either spatially focused pheromone gradients or a change in local plasma membrane curvature initiate this down-regulation. Regardless of the trigger, our findings indicate that such down-regulation requires the cell wall protein Fig2. In the absence of Fig2, mating cells cannot distinguish a contact site from a shmoo tip.

### Mating in the absence of Fig2

Fig2 expression is induced by pheromone and a variety of phenotypes have been described for *fig2* mutants: mating mixtures display hyper-agglutination and improved mating under turbulent conditions, but reduced mating under non-turbulent conditions, especially at cold temperatures (Erdman et al., 1998). Reduced mating is due to reduced fusion and increased lysis, and the zygotes that do form are unusual in having narrower connecting bridges. *fig2* zygotes also show defects in subsequent nuclear fusion and spindle orientation (Erdman et al., 1998; Zhang et al., 2002). Our live-cell imaging of mating *fig2* mutants revealed further unexpected defects, including partial/transient fusion and partial/transient lysis events.

We suggest that a unifying cause for the varied *fig2* defects (except hyper-agglutination, discussed below) is the failure to downregulate the CWI at the contact site. We found that Pkc1 localization was more prominent at the contact site of *fig2* partners, and that cell wall degradation at the contact site was delayed, blocked, or reversed in *fig2* matings. By attempting to repair the cell walls as they are being degraded, mutants would find it harder to successfully remove the walls for fusion. Even when small holes in the cell wall allow fusion to proceed, cell wall repair could close off the fusion pore, yielding transient and unproductive fusion. And even if fusion is not reversed, cell wall repair may make it harder to expand the fusion bridge. A narrower bridge would make it harder for the nuclei to fuse, and diploid nuclei that did fuse might be deformed within the narrow bridge in a manner that causes later defects in zygote spindle orientation.

### How does Fig2 function to enable CWI down-regulation at contact sites?

It has been proposed that like the agglutinins, Fig2 contributes to mating by physically adhering the cell walls of the mating partners (Guo et al., 2000; Huang et al., 2009). Consistent with that view, cells lacking both Fig2 and agglutinins fail to mate (Guo et al., 2000). However, several other observations seem inconsistent with an adhesive role for Fig2. As mentioned above, cells lacking Fig2 hyper-agglutinate, suggesting that if anything, Fig2 acts to reduce cell-cell interaction. Given its large size (1609 residues), a potential explanation for that phenotype is that Fig2 protrudes beyond the surface of the cell wall and reduces the potential for agglutinin interactions (Dranginis et al., 2007). Moreover, unlike the mating-type-specific agglutinins, Fig2 is expressed in both mating types (Erdman et al., 1998), and it is not clear why such a protein would mediate interaction between mating partners but not between cells of the same mating type. In addition, we found that cells lacking Fig2 displayed a constellation of mating defects despite retaining strong attachment between mating types via the agglutinins. Conversely, cells lacking agglutinins showed no mating defect on solid media, despite lacking any strong interaction between partners. Together, these observations suggest that Fig2 plays a distinct role to that of the agglutinins.

We suggest that instead of simply adhering the walls of two mating partners, Fig2’s primary role is to down-regulate the CWI at the contact site. One possibility is that Fig2 is needed to translate a contact-site-specific cue such as local membrane curvature (Smith et al., 2017) into CWI down-regulation. Alternatively, Fig2 may be directly involved in differentiating a contact site from a shmoo tip, although as Fig2 is present at shmoo tips as well as contact sites it is not clear how such a distinction might be achieved. CWI downregulation then enables a tighter focusing of hydrolase secretion that successfully degrades the intervening cell walls to allow fusion.

If *fig2* mutants are unable to detect the difference between a contact site and a shmoo tip, then why do some of them still succeed in fusing? We speculate that a contact geometry-dependent increase in hydrolase activity between partner cells (Huberman and Murray, 2014) may suffice for some wall degradation even without the contact-mediated change in CWI activity. For this mechanism to lead to fusion, the walls of partner cells must remain stably oriented in the same geometry. That stability is provided by the engagement of mating agglutinins at the contact site. In the absence of this tight connection, mating partners simply push each other away and rotate, explaining the complete mating failure of cells lacking both Fig2 and agglutinins.

Our findings leave a key mystery to be addressed: how does a purely extracellular protein like Fig2, with no transmembrane domain, communicate with CWI components in the cytoplasm? One possibility is that Fig2 interacts with the extracellular domains of CWI sensors. Alternatively, Fig2 interactions within the cell walls may affect physical properties of the walls that indirectly regulate the CWI. In either case, it will be fascinating to understand how cell-cell contact between partners is detected and conveyed to intracellular CWI components for successful mating.

## Abbreviations

CWI: Cell Wall Integrity Pathway
GAP: GTPase Activating Protein
GEF: Guanine nucleotide Exchange Factor
MAPK: Mitogen-Activated Protein Kinase
WT: Wild-Type

## ACKNOWLEDGEMENTS

We thank Rossie Clark-Cotton for key observations on *fig2* mutants, Analeigha Colarusso for help with strain construction, and Rachel K Meade for coding in R expertise. For imaging conducted with the aid of the Duke Light Microscopy Core Facility, and we thank Lisa Cameron and Yasheng Gao for assistance. We thank Masayuki Onishi (Duke University), Stephen Bell (MIT), and members of the Lew lab for comments on the manuscript. This work was funded by NIH/NIGMS grant R35GM122488 to D.J.L. The authors declare no competing financial interests.

## MATERIALS AND METHODS

### Yeast Strains

All yeast strains (Table 1) were created in a YEF473 background (*his3-Δ200 leu2-Δ1 lys2-801 trp1-Δ63 ura3-52*) (Bi and Pringle, 1996) and were constructed using standard molecular genetic techniques.

| Strain | Relevant Genotype | Source |
| --- | --- | --- |
| DLY8156 | MATα | This Study |
| DLY9069 | MATa BEM1-GFP:LEU2 | (Kozubowski et al., 2008) |
| DLY12944 | MATα BEM1-tdTomato:HIS3 | (Howell et al., 2012) |
| DLY23438 | MATα BEM1-GFP:LEU2 fig2::HYG<sup>R</sup> aga1::KAN<sup>R</sup> | This Study |
| DLY23441 | MATa BEM1-tdTomato:HIS3 fig2::HYG<sup>R</sup> aga1::KAN<sup>R</sup> | This Study |
| DLY23471 | MATα BEM1-tdTomato:HIS3 sag1::NAT<sup>R</sup> | This Study |
| DLY23477 | MATa BEM1-GFP:LEU2 aga2::KAN<sup>R</sup> | This Study |
| DLY24009 | MATa BEM1-GFP:LEU2 PKC1-TdTomato:HIS3 | This Study |
| DLY24084 | MATa BEM1-GFP:LEU2 PKC1-TdTomato:HIS3 fig2::HYG<sup>R</sup> aga1::KAN<sup>R</sup> | This Study |
| DLY24254 | MATa BEM1-tdTomato:HIS3 WSC1-GFP:KAN<sup>R</sup> | This Study |
| DLY24325 | MATa BEM1-tdTomato:HIS3 ROM2-mNG:NAT<sup>R</sup> | This Study |
| DLY24546 | MATa fig2::HYG<sup>R</sup> | This Study |
| DLY24551 | MATα BEM1-GFP:LEU2 PKC1-TdTomato:HIS3 fig2::HYG<sup>R</sup> | This Study |
| DLY24655 | MATα BEM1-tdTomato:HIS3 PKC1-mNG:NAT<sup>R</sup> sag1::NAT<sup>R</sup> | This Study |
| DLY24657 | MATa aga2::KAN<sup>R</sup> | This Study |
| DLY24672 | MATa GFP:LEU2 fig2::HYG<sup>R</sup> | This Study |
| DLY25318 | MATα Psr1(1-28aa)-mCherry:LEU2 HTB2-mCherry:HIS3 fig2::HYG<sup>R</sup> aga1::KAN<sup>R</sup> | This Study |
| DLY25319 | MATa GFP:LEU2 HTB2-mCherry:HIS3 fig2::HYG<sup>R</sup> | This Study |
| DLY25320 | MATα GFP:LEU2 HTB2-mCherry:HIS3 | This Study |
| DLY25321 | MATα Psr1(1-28aa)-mCherry:LEU2 HTB2-mCherry:HIS3 fig2::HYG<sup>R</sup> | This Study |
| DLY25323 | MATa GFP:LEU2 HTB2-mCherry:HIS3 | This Study |
| DLY25324 | MATα Psr1(1-28aa)-mCherry:LEU2 HTB2-mCherry:HIS3 | This Study |
| DLY26219 | MATα Psr1(1-28aa)-mCherry:LEU2 fig2::HYG<sup>R</sup> | This Study |

The following alleles have been previously described: *BEM1*-GFP:*LEU2* (Kozubowski et al., 2008), *BEM1*-tdTomato:*HIS3* (Howell et al., 2012), and *PKC1*-tdTomato:*HIS3* (Lai et al., 2018).

*WSC1*-GFP:kan^r^ and ROM2-mNeonGreen:nat^r^ were constructed using the one-step PCR method (Longtine et al., 1998) in which the fluorophore open reading frames were amplified off a template with 50 BP of homology to the C terminus and 3’ UTR region to promote homologous recombination at the native locus. Templates used were pFA6a-GFP(S65T)-KanMX6 (Bähler et al., 1998) and *pNCS-mNeonGreen:nat^R^*(Allele Biotechnology). Correct integration was confirmed by PCR.

To delete *AGA1*, *AGA2*, *FIG2*, and *SAG1* we used a PCR based method (Baudin et al., 1993) to replace the open reading frame with a selectable marker. Primers with 50 bp of homology to the 5’ and 3’ region of the gene to be deleted were used to amplify the selectable marker from a template. Templates used were pFA6a-kanMX6 (Longtine et al., 1998), pRS40H-hphMX4 and pRS40N-natMX4 (Chee and Haase, 2012), pRS314 (Sikorski and Hieter, 1989). Correct integration was confirmed by PCR using primers designed to amplify the flanking region.

### Live Cell Microscopy

Cells were grown overnight at 30 °C in Complete Synthetic Medium (CSM; MP Biomedicals) with 2% dextrose (CSM+Dex) to mid-log phase (10^6^–10^7^ cells/ml). For mating assays, a total of 6^6^ cells were spun down in a 1:1 ratio of each mating type at 12,000 RPM for 1 min. Cells were resuspended in 3.6 μL of CSM+Dex and.6 μL of the resuspended cells (∼10^6^ cells) were plated on agarose pads. Agarose pads were prepared by sandwiching 200 μL CSM with 2% dextrose and 2% agarose between microscope slide and coverslip. For shmooing experiments, 10 μM of alpha factor were added to agarose pads during preparation. All slides were sealed with petroleum jelly for the duration of the experiment. Time lapse movies were acquired at 30°C in a humidity controlled chamber at 2 min intervals. The entire cell volume was acquired using 15 z-stacks spaced.4 um apart. Imaging was performed using two microscopes:

i. an Andor XD Revolution inverted microscope with a CsuX-1 5000 rpm spinning-disk head (Yokogawa) controlled by MetaMorph 7.8 ID 8562. Images were captured using an iXon 3 897 EMCCD camera and then a subsequent iXon Life 888 EMCCD. A 100x/1.4 oil U PlanSApo DIC, WD: 0.13 mm, ∞/0.17/FN26.5, UIS2 objective was used. A laser power of 13% and exposure time of 250 ms was used for both the 488 nm green fluorescence channel and the 561 nm red fluorescence channel. Images were denoised using the ImageJ Hybrid 3D Median Filter Plugin (2007) with drift correction completed using HyperStackReg plugin in Fiji, written by Ved Sharma [DOI.10.5281]. All images shown and analyzed are maximum intensity projections.
ii. a Nikon Ti2E inverted microscope with a CSU-W1 spinning-disk head (Yokogawa) controlled by NIS-Elements software (Nikon Instruments). Images were captured using a Hamamatsu ORCA Quest qCMOS camera. A CFI60 Plan Apochromat Lambda D 100x Oil Immersion Objective Lens, N.A. 1.45, W.D. 0.13 mm, F.O.V. 25 mm, DIC, Spring Loaded was used. A laser power of 180 μW and exposure time of 175 ms was used for the 488 nm green fluorescence channel. A laser power of 125 μW and exposure time of 175ms was used for the 561 nm red fluorescence channel. Images were denoised using the median filter in NIS-Elements General Analysis 3 (GA3, Nikon Instruments) software drift correction completed using HyperStackReg plugin in Fiji, written by Ved Sharma [DOI.10.5281]. All images shown and analyzed are maximum intensity projections.

### Image Analysis

#### Mating Efficiency

To score mating efficiency only cells that had an opportunity to mate were analyzed. Cells with the opportunity to mate were defined as cells that were touching a cell of the opposite mating type that was in G1 phase. Cells that only had an opportunity to mate for less that 20 min before the movie ended were excluded from analysis. Cells whose potential mating partner instead mated with a different cell were excluded from analysis. Cells with the opportunity to mate were scored as mated or not mated based on visual mixing of Bem1 cytoplasmic signal that occurs at the time of fusion.

#### Clustering Parameter

To quantify the degree of protein clustering for Bem1, Wsc1, Rom2, and Pkc1, a deviation from uniformity metric (referred to as Clustering Parameter (CP)) was calculated from a maximum intensity projection. An elliptical Region of Interest (ROI) was drawn around each cell for each time point measured. For cells undergoing mating, the 30 min leading up until fusion were measured. For shmooing cells, measurements were obtained for every timepoint throughout the 3 hour movie. CP was measured using a MATLAB based GUI called ROI_TOI_QUANT_V9, developed by Denis Tsygankov (Lai et al., 2018).

#### Linescans

Linescans shown were collected in Fiji. Using the Freehand Line Tool, the fluorescent intensities around the perimeter of the cell were obtained by averaging the values in a 3-5 pixel wide line (microscope depending). Measurements were stored within the ROI Manager for each timepoint measured and exported for visualization within R ggplot. Linescans collected were either: i) the time point 2 min before fusion. ii) a sum projection in time of the 30 min in 2 min intervals leading up until fusion.

#### Kymographs

To generate kymographs, linescans were first collected in Fiji for the 30 min leading up until fusion using the above method. For mating crosses that experienced fusion, linescans were obtained at 2 min intervals for the 30 min leading up to fusion. Fusion was excluded from analysis. For mating crosses where fusion was abolished, linescans were obtained at 2 min intervals for the 3 h duration of the movie. For shmooing experiments, linescans were obtained at 2 min intervals for the 3 h duration of the movie. Measurements were stored within the ROI Manager for each timepoint measured and exported using the Multi Plot function for further analysis and data visualization within R. Linescans were aligned such that the maximum intensity of the Bem1 signal 2 min prior to fusion is positioned at the center of the graph and designated by a dotted red line. Positioning of prior Bem1 signal and CWI Pathway signal is all shifted in relation. Individual kymographs were generated in R using ggplot.

## Statistical Analysis

All statistics were completed in R using the R Stats package. Kolmogorov-Smirnov Tests were conducted using ks.test. Chi Squared tests were conducted using chisq.test.

## Data Availability

The data generated in this study are available from the corresponding author upon reasonable request.

## Supplemental Material

Supplemental Table 1 provides a list of all strains used in this study.

